# ACAT1/SOAT1 Blockade Suppresses LPS-Mediated Neuroinflammation by Modulating the Fate of Toll-Like Receptor 4 in Microglia

**DOI:** 10.1101/2022.08.30.505911

**Authors:** Haibo Li, Thao N. Huynh, Michael Tran Duong, James G. Gow, Catherine C.Y. Chang, Ta-Yuan Chang

**Affiliations:** Department of Biochemistry and Cell Biology, Geisel School of Medicine at Dartmouth, Hanover, NH 03755 USA; Department of Bioengineering, Perelman School of Medicine, University of Pennsylvania, Philadelphia, Pennsylvania, USA

**Keywords:** Cholesterol, cholesteryl esters, acyl-CoA:cholesterol acyltransferase, Sterol O-acyltransferase, ACAT inhibitor, lipid rafts, toll-like receptor 4, neuroinflammation, Alzheimer’s disease

## Abstract

**Background:** Cholesterol is essential for growth and maintenance of mammalian cells. It is stored as cholesteryl esters by the enzymes acyl-CoA:cholesterol acyltransferases 1 & 2 (ACAT 1 & 2) (Sterol O-acyltransferase 1 & 2; SOATs in GenBank). ACAT1 blockade (A1B) in macrophages ameliorates various pro-inflammatory responses elicited by lipopolysaccharides (LPS) or by cholesterol loading. In mouse and human brains, *Acat1* expression dominates over *Acat2* and *Acat1* is elevated in many neurodegenerative diseases and in acute neuroinflammation. However, the possible effects of ACAT1 blockade in neuroinflammation, regulated by mediators such as Toll-Like Receptor 4 (TLR4), has not been studied.

**Methods:** We conducted LPS-induced acute neuroinflammation experiments in control vs myeloid specific or neuron specific *Acat1* knockout (KO) mice. Furthermore, we evaluated LPS-induced neuroinflammation in the microglial cell line N9 with or without pre-treatment of the small molecule ACAT1-specific inhibitor K-604. Biochemical and microscopy assays were used to monitor inflammatory responses and the fate of TLR4.

**Results:** *In vivo* studies revealed that *Acat1* inactivation in myeloid cell lineage, but not in neurons, markedly attenuated LPS-induced activation of various pro-inflammatory response genes in hippocampus and cortex. Studies in cell culture showed that pre-incubating cells with K-604 significantly ameliorated the pro-inflammatory responses induced by LPS. In cells acutely treated with LPS (for 30 min), pre-incubation with K-604 significantly increased the endocytosis of TLR4, the major transmembrane signaling receptor that mediates LPS-dependent acute neuroinflammation. In cells chronically treated with LPS (for 24-48 hrs), pre-incubation with K-604 significantly decreased the total TLR4 protein content, presumably due to enhanced trafficking of TLR4 to the lysosomes for degradation. For *ex vivo* evidence, we isolated microglia from adult mice, and found that in mice without LPS stimulation, myeloid *Acat1* inactivation altered cellular distribution of TLR4; in mice with LPS stimulation, myeloid *Acat1* inactivation decreased the cellular content of TLR4.

**Conclusion:** Blocking ACAT1 in mouse microglia alters the fate of TLR4 and suppresses its ability to participate in pro-inflammatory signaling cascade in response to LPS.

## Introduction

Cholesterol is an essential lipid for the growth and maintenance of all mammalian cells. Its metabolites include oxysterols, neurosteroids, bile acids, and steroid hormones; these substances also play important physiological functions [1,2]. The brain is the most cholesterol rich organ in the body; while it only constitutes only 2.1% of total body weight, it contains 23% of the body’s total cholesterol [3,4]. Increasing evidence indicates that cholesterol dyshomeostasis is closely linked with several neurodegenerative diseases, including Alzheimer’s disease (AD) [1,2]. Excess (free, unesterified) cholesterol is stored as cholesterol esters (CEs). Normally, CE levels in mice and human brains are very low. In contrast, in the vulnerable regions of brain samples (such as entorhinal cortex) from late onset Alzheimer’s disease (LOAD) patients, CE levels were found to increase by 1.8-fold [1]. In addition, in the relevant brain regions of three different AD mouse models, the CE levels were also found to be 3- to 11-fold higher than those in controls [6,7]. These findings suggest that CE content positively correlates with AD risk. What causes CEs to be elevated in the AD brains is an active area of research. For biosynthesis of CEs, there are two distinct genes, acyl-CoA:cholesterol acyltransferase 1 (*Acat1*) and 2 (*Acat2*) (also called sterol O-acyltransferase 1 and 2 (*Soat1,2*) in GenBank), encoding two distinct enzymes, ACAT1 [2] and ACAT2 [9-11]. Both enzymes use long-chain fatty acyl-CoAs, and sterols with 3-beta–OH, including cholesterol and various oxysterols, as their substrates [3]. ACAT1 is ubiquitously expressed in essentially all cells, including cells in peripheral tissues and in the brain; ACAT2 is mainly expressed in intestinal enterocytes and in hepatocytes; low levels of ACAT2 are also detectable in various peripheral tissues [4]. Both ACAT1 and ACAT2 are integral membrane proteins located in the ER region, and both are allosterically activated by cholesterol or oxysterols, as reviewed in [5]. CEs are part of lipid droplets; they cannot substitute the functions of cholesterol.

Recent evidence from several laboratories cast new light on ACAT1 as a promising molecular target for the treatment of AD. AD pathological hallmarks consist of extracellular amyloid plaques, composed of amyloid beta peptides (Aβ; especially Aβ1-42), and neurofibrillary tangles (composed of hyperphosphorylated tau). Mechanistically, ACAT1 blockade (A1B) acts by (**1**) reducing CE and amyloid pathology [15-17]; (**2**) increasing the content of the neuroprotective oxysterol 24(S)-hydroxycholesterol in the 3XTg AD mouse brain [6] and in the AD patient induced pluripotent stem cells (iPSC) derived from human neurons [7]; (**3**) increasing autophagy flux leading to the clearance of Aβ oligomers in microglia [8], and the clearance of misfolded tau in neurons [9]; (**4**) preventing the inhibitory effects of CEs on hyperphosphorylated tau degradation by proteostasis in EOAD patient-derived neurons [7]; (**5**) decreasing the protein content of mutant full-length human amyloid precursor protein (hAPP) in the brains of the 3XTg AD model [6], and in AD patient iPSC derived human neurons [7]; (**6**) clearing CEs accumulated in myelin debris treated microglia that lack TREM-2, a risk factor for LOAD [10]. These results show that in various AD models, A1B offers benefits to suppress amyloidopathy and tauopathy. On the other hand, LOAD is also accompanied by chronic neuroinflammation [11].

To our knowledge, whether A1B affects neuroinflammation in AD models has not been reported. We had previously reported that, in a mouse model, A1B by genetic inactivation of *Acat1* in the myeloid lineage reduces inflammatory responses in adipose tissue macrophages and protects against diet induced obesity [12]. In addition, in a different mouse model, myeloid *Acat1* KO attenuates pro-inflammatory responses in macrophages and protects against atherosclerosis [13]. Macrophages and microglia belong to the same myeloid cell lineage. In the central nervous system (CNS), microglia play a key role in mediating neuroinflammation [14]. Based on our previous work in macrophages, we hypothesize that A1B may also suppress pro-inflammatory responses in microglia. Inflammation maybe acute or chronic. At the mechanistic level, acute inflammation and chronic inflammation share many commonalities. It is known that a single, systemic administration of lipopolysaccharides (LPS) produces acute neuroinflammation in adult mice through transcriptional pathways activated by classic immune regulators including TLR4 [15]. In the current work, we use LPS-induced acute neuroinflammation as the model to test our hypothesis and report our findings.

## Methods

### Animals and acute neuroinflammation induction

WT, *Acat1^flox/flox^*, *Acat1^flox/flox^ LysM^Cre^* (myeloid cell-specific *Acat1* knockout), *Acat1^flox/flox^Syn1^Cre^* (neuronal cell-specific *Acat1* knockout) mice were all in the C57BL/6J background. The *Acat1^flox/flox^* and *Acat1^flox/flox^LysM^Cre^* mice were generated by crossing *Acat1^flox/flox^* mice with *LyzM^Cre^* mice (The Jackson Laboratory) as previous described [16]. *Acat1^flox/flox^Syn1^Cre^* mice were generated by crossing *Acat1^flox/flox^* mice with *Syn1^Cre^* mice (The Jackson Laboratory). Littermates produced from relevant mice were used to conduct experiments as described in **Figs 1** and **2**. Mice were housed in a specific pathogen-free barrier facility under a regular light-dark cycle and fed standard chow. All mouse protocols were approved by the Dartmouth College Institutional Animal Care and Use Committee and followed NIH guidelines. Acute neuroinflammation was induced by intraperitoneal lipopolysaccharide (LPS) injection of 5 mg/kg body weight as previous described [26, 27]. LPS was obtained from Santa Cruz (sc-3535) and dissolved in sterile PBS as a 5 mg/ml stock solution. 24 hrs later, mice were perfused with sterile HBSS (Corning21-022-CV) and sacrificed. Hippocampus and cerebral cortex were isolated and homogenized for RNA extraction.

**Fig 1.**
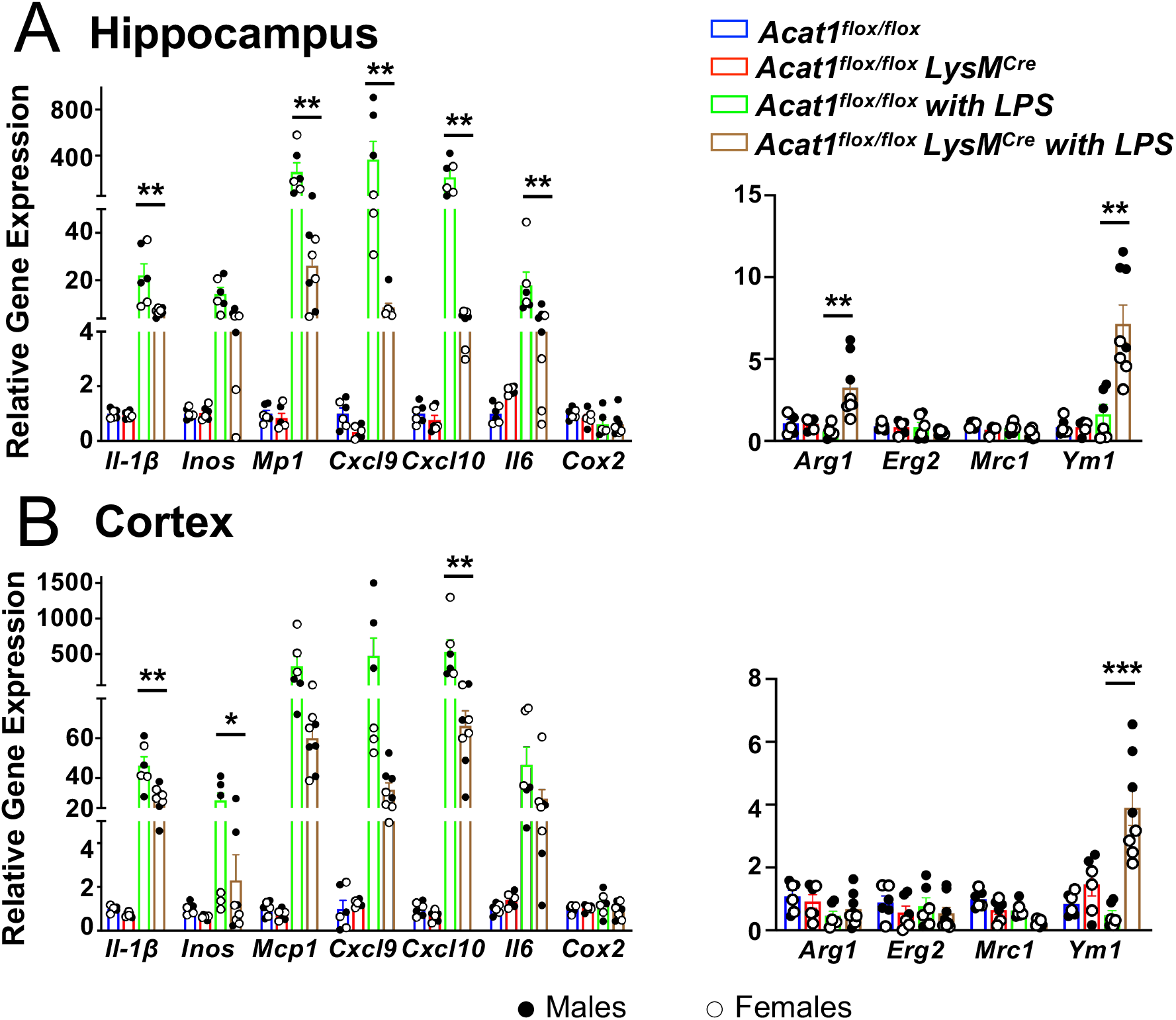
Myeloid *Acat1* blockade alters the inflammatory gene profile in hippocampus (A) and cortex (B) after LPS injection. Two-month-old *Acat1^flox/flox^* and *Acat1^flox/flox^ LyzM^Cre^* (myeloid Acat1 knockout) mice as littermates were injected peritoneally with LPS at 5 mg/kg body weight. 24 hrs later, mice were sacrificed, and mRNA was extracted from hippocampus and cortex. qPCR was performed for pro-inflammatory (left panel) or anti-inflammatory (right panel) gene expression. ● indicates male mice and ○ indicates female mice.

**Fig 2.**
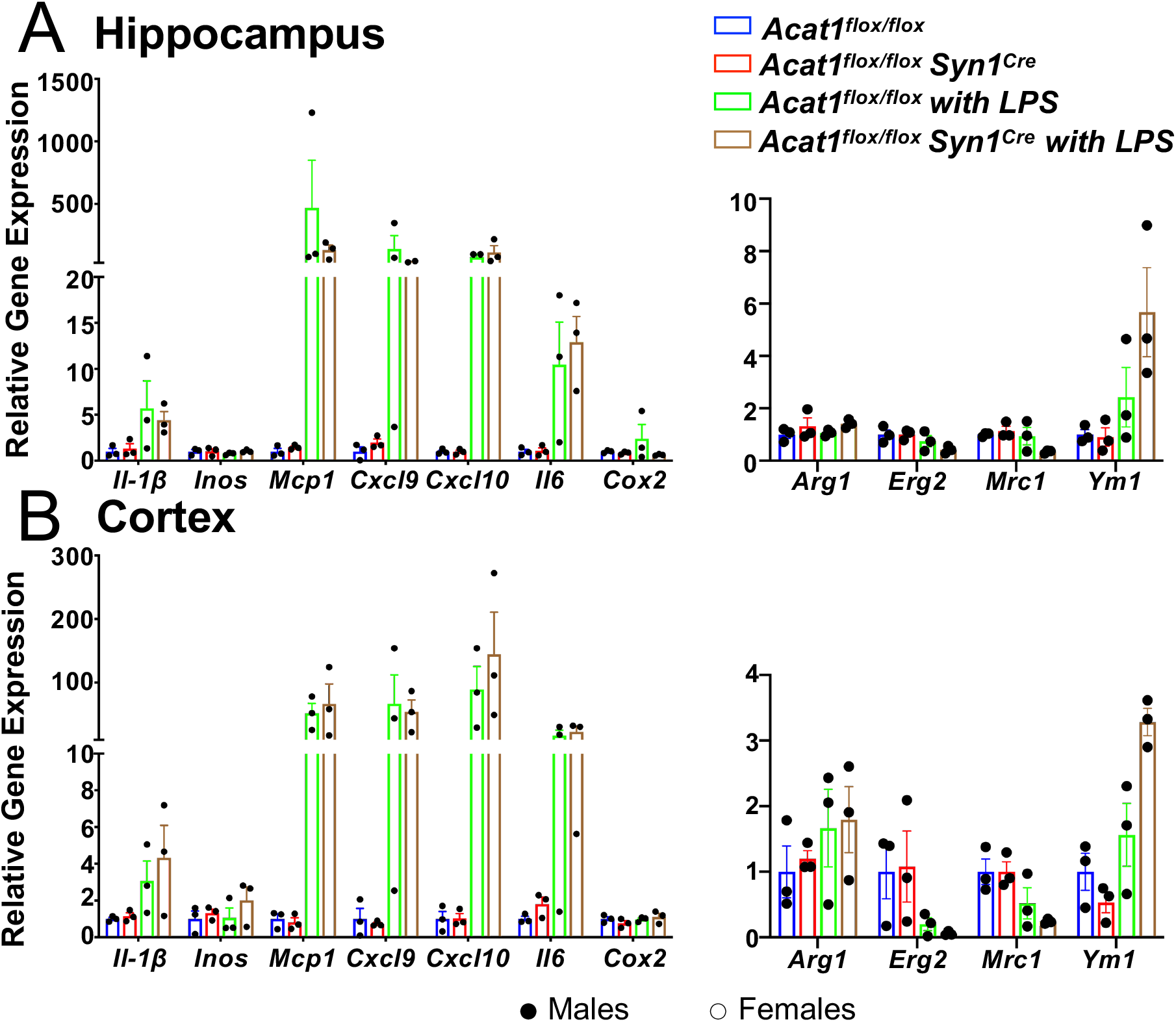
Neuron-specific *Acat1* blockade did not change the inflammatory gene expression in hippocampus and cortex from after LPS injection. Two-month-old *Acat1 ^flox/flox^* mice and *Acat1 ^flox/flox^ Syn1^Cre^*(neuronal cell specific *Acat1* knockout) mice as littermates were injected peritoneally with LPS at 5 mg/kg body weight. 24 hrs later, mice were sacrificed, and mRNA was extracted from hippocampus and cortex. qPCR was performed for pro-inflammatory (left panel) or anti-inflammatory(right panel) gene expression. ● indicates male mice and ○ indicates female mice.

### Cell culture

Mouse N9 microglial cells were maintained in RPMI-1640 with 10% calf serum at 37°C with 5% CO_2_ in a humidified incubator as previously described [8]. For ACAT1 inhibition, ACAT1 inhibitor K-604 was first dissolved in DMSO at 5 mM as stock solution and diluted into the culture medium such that the final concentration was at 0.5 μM as previously described [8].

### RNA Isolation and qPCR

The procedure was as described previously [12]. Total RNAs were isolated using TRIzol reagent (Invitrogen). 2.5 μg of total RNA was treated with DNase I (New England BioLabs M0303) to remove any remaining genomic DNA. 0.5 μg DNase I treated RNA was reverse-transcribed using iScript cDNA Synthesis Kit (Bio-Rad) to prepare cDNA. qPCR was performed using iTaq Universal SYBR Green Supermix (Bio-Rad). The following cycles were performed: an initial denaturation cycle of 94°C for 5 min, followed by 40 amplification cycles of 94°C for 15 s and 60°C for 1 min. Relative quantification was determined using ΔΔCT method. The mRNA expression values were normalized with β-actin mRNA level. The following primers were used:

***β-actin*** forward: 5′-CAACGAGCGGTTCCGAT-3′, reverse: 5′-GCCACAGGATTCCATACCCA-3′;

***il-1β*** forward: 5′- ACAGAATATCAACCAACAAGTGATATT-3′, reverse: 5′-GATTCTTTCCTTTGAGGCCCA-3′;

***inos*** forward: 5′-ACATCGACCCGTCCACAGTAT-3′, reverse: 5′-CAGAGGGGTAGGCTTGTCTC-3′;

***mcp1*** forward: 5′- CCCACTCACCTGCTGCTACT-3′, reverse: 5′-TCTGGACCCATTCCTTCTTG-3′;

***cxcl9*** forward: 5′- GGAGTTCGAGGAACCCTAGTG-3’, reverse: 5′-GGGATTTGTAGTGGATCGTGC-3′;

***cxcl10*** forward: 5′- CCAAGTGCTGCCGTCATTTTC-3′, reverse: 5′-GGCTCGCAGGGATGATTTCAA-3′;

***il6*** forward: 5′-TCCAGTTGCCTTCTTGGGAC-3′, reverse: 5′- GTACTCCAGAAGACCAGAGG-3′;

***cox2*** forward: 5′- AGCAACCCGGCCAGCAATCT-3′, reverse: 5′-CCTGCTGCCCGACACCTTCA-3′;

***arg1*** forward: 5′- CTCCAAGCCAAAGTCCTTAGAG-3′, reverse: 5′-AGGAGCTGTCATTAGGGACATC-3′;

***erg2*** forward: 5′- GCAAAGGACCTTGATGGAGC-3′, reverse: 5′-GGCCTAAGTTTTCGGAAGGC-3′;

***mrc1*** forward: 5′- CTCTGTTCAGCTATTGGACGC-3′, reverse: 5′-CGGAATTTCTGGGATTCAGCTTC-3′;

***ym1*** forward: 5′- CATGAGCAAGACTTGCGTGAC-3′, reverse: 5′-GGTCCAAACTTCCATCCTCCA-3′

### OptiPrep^™^ subcellular fractionation

Experiments were performed at 4°C as previously described [17]. Briefly, N9 cells were seeded onto 150 mm plates at a density of 10,000 cells/mL (200,000 cells/plate). 68 and 72 hrs later, 0.5 μM K-604 (or DMSO as control) and 200 ng/mL LPS (or PBS as control) were added into the plates respectively. 30 min after adding LPS, cells were scraped off into 1 mL homogenization buffer containing 0.25 M sucrose, 20 mM Tris buffer (pH 7.4), 1 mM EDTA and protease inhibitor cocktail (Sigma) and were homogenized using a stainless-steel tissue grinder (Dura-Grind, Wheaton). The post-nuclear supernatant was loaded onto the top of a 9 mL, continuous 5–20% OptiPrep^™^ gradient in homogenization buffer and was fractionated by sedimentation velocity ultracentrifugation in a SW41 rotor at 40,000 rpm for 3 hrs; 15 equal fractions were collected from top to bottom for Western Blot analyses.

### Whole cell protein isolation and western blot analyses

For whole cell protein isolation, cells were harvested in RIPA buffer containing protease inhibitor cocktail (Sigma) or with phosphatase inhibitor cocktail (Sigma) for phosphorylated proteins. Followed by sonication and centrifugation, the protein concentration of the supernatant was determined by Lowry protein assay. The lysates were run on a 10% SDS-PAGE gel and transferred to a 0.45 μm nitrocellulose membrane for 4 hrs at 300 mA. After blocking in 5% milk in TBST buffer for 1 hr at room temp, the membranes were incubated with anti-TLR4 (Santa Cruz sc-293072), anti-p-IκB-α(Santa Cruz sc-8404), anti-IκB-α (Santa Cruz sc-1643), anti-Na+/K+ATPase α1 (Santa Cruz sc-21712), anti-LAMP1(Santa Cruz sc-20011), or anti-PMCA (Plasma membrane-type Ca^2+^-ATPases) (Santa Cruz sc-271917), along with anti-vinculin (Novus NB600-1293) antibodies, or anti-β-Actin Antibody(Santa Cruz sc-47778) as protein loading control.

### TLR4 endocytosis assay

TLR4 endocytosis was assayed as described previously [18]. In brief, N9 cells were seeded on poly-d-lysine-coated glass cover slides in 12-well plates at a density of 150,000 cells/well and went through different treatments as indicated in **Fig 5** legend. Cells were incubated in RPMI (with 5% goat serum) with 1:200 dilution of specific anti-TLR4 (Santa Cruz sc-293072) antibodies at 4°C for 1 hr then washed twice in cold RPMI to remove unbound antibodies. Subsequently, cells were re-incubated in RPMI containing 10% calf serum at 37°C in cell incubator for 1 hr. To remove cell surface-bound antibodies, the cells were washed twice for several seconds in RPMI adjusted to pH 2.0. Cells were then fixed with 4% paraformaldehyde for 10 min at 4°C, permeabilized with 0.3% Triton for 20 min at 37°C, stained with goat anti-mouse secondary antibodies coupled to Alexa Fluor 568 for 1 hr at 37°C. Finally, the coverslips were mounted on microslides with a drop of ProLong Gold antifade reagent with DAPI (Invitrogen) for imaging under a confocal fluorescence microscopy.

### Immunofluorescence microscopy

Cells were grown overnight on poly-d-lysine-coated glass cover slides in 12 well plates at a density of 150,000–300,000 cells/well. Cells were washed with PBS and fixed with 4% paraformaldehyde for 10 min at 4°C, then washed cells three times with PBS, and permeabilized with 0.1% or 0.3% Triton X-100 in PBS for 10-20 min followed by PBS washes three times. Then the slides were blocked for 1 hr at room temp with 5% goat serum in PBS, incubated overnight at 4°C with anti-TLR4 (Cell Signaling 14358, Invitrogen #PA5-32124), anti-LAMP1(Santa Cruz sc-20011), or anti-N-cadherin (Invitrogen MA1-91128) antibodies in blocking buffer, then washed three times with PBS and incubated with AlexaFluor dye-conjugated secondary antibodies for 1 hr at room temp. Afterwards, cells were washed three times with PBS, rinsed with double-distilled water, and mounted on glass slides with a drop of ProLong Gold antifade reagent with DAPI (Invitrogen). Confocal images were obtained by using a Zeiss LSM 510 confocal microscope. Image analysis was performed using ImageJ software.

### Statistical analysis

All statistical analysis was performed using Prism8 software (GraphPad). A two-tailed Student’s t test was used when two values were compared. For multiple comparisons, a one-way ANOVA with a Tukey’s post-hoc test was used. Error bars indicate SEM. * P<0.05; ** P<0.01; *** P<0.001; ****P<0.0001.

## Results

### *ACAT1* gene expression is elevated in neurodegenerative diseases and in acute neuroinflammation

In humans and mice, there are two ACAT/SOAT genes. To document the relative expression of ACAT/SOAT in various cell types in the brain, we first retrieved transcriptomics data from the Barres Laboratory [19] and assessed expressions of *SOAT1*/*Soat1* and *SOAT2*/*Soat2* in the cortex of human and mouse brains under normal condition (**Fig. S1**). The data indicate that, in both humans and mice, microglia are the main cell types that express *SOAT1*; astrocytes and neurons also express *SOAT1*, but at lesser levels. In various cell types examined, the expression of *SOAT1* mRNA predominates over that of *SOAT2* by at least 10-fold. In the CNS, chronic inflammation produces cytokines that activate microglia. To determine the expression levels of microglial ACAT1 under neuroinflammation condition, we obtained transcriptomics data from Friedman and colleagues [20]. These data described gene expression profiles from purified, CD11b^+^ CNS myeloid cells isolated from mouse models for various diseases/conditions (**Fig. S2**). The data indicated that, in microglia, *Soat1* is significantly elevated in many neurodegenerative diseases, including several AD mouse models (three AβPP models and two Tau models), in two of three acute inflammation/infection mouse models, and in a mouse model for aging (out to 22 months of age). In humans, additional data presented in **Fig. S2** (1^st^ bar in brown color) showed that *SOAT1* is elevated in the disease-associated region of late onset Alzheimer’s disease (LOAD). Based on these data, elevated CEs observed in the vulnerable regions of LOAD, and in mouse models for amyloidopathy [1, 21] can be attributed, at least in part, to elevated *SOAT1/Soat1* expression in microglia located in these regions.

### Blocking ACAT1 in myeloid attenuated LPS-mediated acute neuroinflammation in the CNS

In mouse model with obesity or with atherosclerosis, genetic inactivation of *Acat1/Soat1* in the myeloid lineage reduces inflammatory responses in macrophages [23,24]. Microglia and macrophages belong to the same myeloid lineage. We suspect that blocking ACAT1 in myeloid lineage cells may also reduce inflammatory responses in the CNS. To test this hypothesis, we treated 2-month-old, sex-matched *Acat1^flox/flox^* and *Acat1^flox/flox^ LysM^Cre^* (designated as *Acat1^−M/−M^*) mice as littermates with a single intraperitoneal injection of 5 mg/kg LPS to elicit acute neuroinflammation. An equal volume of PBS was used as the negative control. One day later, we sacrificed the mice and prepared brain homogenates from both hippocampal and cortex regions to perform q-PCR analyses of various genes. Based on our previous work [23,24], the relevant genes that we chose to analyze included *Il1-β, Inos, Mcp1, Cxcl9, Cxcl10, Il6, Cox2* (representing pro-inflammatory genes), as well as *Arg1, Erg2, Mrc1, and Ym1* (representing anti-inflammatory genes). The results (**Fig. 1A**; left panel) show that in the hippocampal region, at the basal state, myeloid A1B did not significantly alter the gene expressions of various pro-inflammatory genes. Injection of LPS highly activated the expressions of multiple pro-inflammatory genes; the fold increases occurred by 20- to 400-fold, in a gene-specific manner. The only exception was *Cox2*; its expression did not respond to LPS. Consistent with our hypothesis, myeloid A1B significantly attenuated the LPS-induced gene expressions of *Il1-β* (by ~70%), *Mcp1* (by ~70%), *Cxcl9* (by ~90%), *Cxcl10* (by ~98%), and *Il6* (by ~98%). In addition, A1B tended to decrease the gene expression of *Inos*, but its effect did not reach statistical significance. Additional results (**Fig 1B**; left panel) show that similar findings occurred in the cortex region: LPS injection elicited large increases in the expressions of various pro-inflammatory genes, and A1B in myeloid lineage significantly attenuated the expressions of these genes. In terms of the expressions of anti-inflammatory genes (**Fig 1A, B**; right panels): at the basal state (without LPS), in both hippocampal and cortex regions, A1B in myeloid did not significantly alter the gene expressions of any of these genes. LPS injection to the *Acat1^flox/flox^* mice did not significantly affect the expressions of these genes; LPS injection to the *Acat1^−M/-M^* mice caused significant increases in the gene expressions of both *Arg1* (by ~7-fold) and *Ym1* (by ~4-fold) in the hippocampal region (**Fig 1A;** right panel); in the cortex region, LPS injection to the *Acat1^−M/-M^* mice elicited a similar stimulating effect on the *Ym1* gene but not the *Arg1 gene* (**Fig 1B**; right panel). These results show that in response to acute inflammation, myeloid A1B attenuated pro-inflammatory responses and increased anti-inflammatory responses in the CNS. In a separate experiment, we performed a similar experiment as described in **Fig. 1**, but by comparing *Acat1^flox/flox^* and *Acat1*^flox/flox^ *Syn1^Cre^* (neuronal cell specific *Acat1* knockout) to examine the effects of A1B in neurons on various pro-inflammatory and anti-inflammatory genes in response to LPS injection. The results (**Fig. 2**) show that while LPS injection elicited profound increases in various pro-inflammatory genes in both hippocampus and in cortex; unlike A1B in myeloid, A1B in neurons did not significantly alter the inflammatory gene expression patterns caused by LPS injection. These results are consistent with the notion that in the CNS, LPS-mediated acute inflammation occurs mainly via microglia.

### Blocking ACAT1 with the small molecule ACAT1 inhibitor K-604 attenuated the pro-inflammatory responses in LPS treated microglial cell line N9

We next aimed at elucidating the action of A1B in response to acute inflammation in microglial cell culture. The mouse microglial cell line N9 closely mimics the phenotypes of mouse primary microglia. For instance, in terms of studying the effects of A1B on autophagy and on cellular metabolism, we had previously shown that the results obtained in N9 cells could all be replicated in the primary microglia isolated from mouse brains [8]. Other laboratories have also reported that N9 is a suitable cell line with which to investigate the inflammatory response towards LPS treatment [34,35]. N9 cells grown with 10% calf serum were pre-treated for 4 hrs with DMSO (control group) or 0.5 μM K-604 [22], then treated with 10, 100, or 1000 ng/mL of LPS for 6 hrs. Afterwards, mRNA was extracted, and qPCR was performed to monitor the expressions of seven pro-inflammatory (**A**) or three anti-inflammatory (**B**) genes. The results (**Fig 3A**) show that the expressions of 5 of the 7 pro-inflammatory genes (*Il1-β, Inos, Cxcl9, Cxcl10*, and *Il6*); dramatically increased (by 10- to over 100-fold, in a gene specific manner, in response to the three doses of LPS tested. The results also show that, in all 5 genes, pre-treating cells with A1B for 4 hrs significantly attenuated their increased expressions (by 25 to 80%, in a gene-specific manner), in response to LPS. Additional results (**Fig 3B**) show that the expression of one of the anti-inflammatory genes tested (*Arg1*) is actually significantly increased in response to LPS (by around 5-fold); pre-treating cells with A1B did not affect the action of LPS. The expressions of the other two anti-inflammatory genes (*Erg2* and *Mrc1*) tended to decrease in response to LPS, but the difference did not reach significance; pretreating cells with A1B did not affect the action of LPS on *Erg2* or on *Mrc1*. The expression of *Ym1* was not calculated due to very low expression (data not shown). Together, these results show that in N9 cells, the pro-inflammatory responses towards LPS mimic those characterized in the mouse CNS; in addition, treating N9 cells with K-604 attenuated the pro-inflammatory responses elicited by LPS, mimicking the anti-inflammatory actions of A1B by myeloid specific *Acat1* KO in the CNS of mice *in vivo*. These results support the use of N9 cells as a cell culture model to elucidate the actions of A1B.

**Fig 3.**
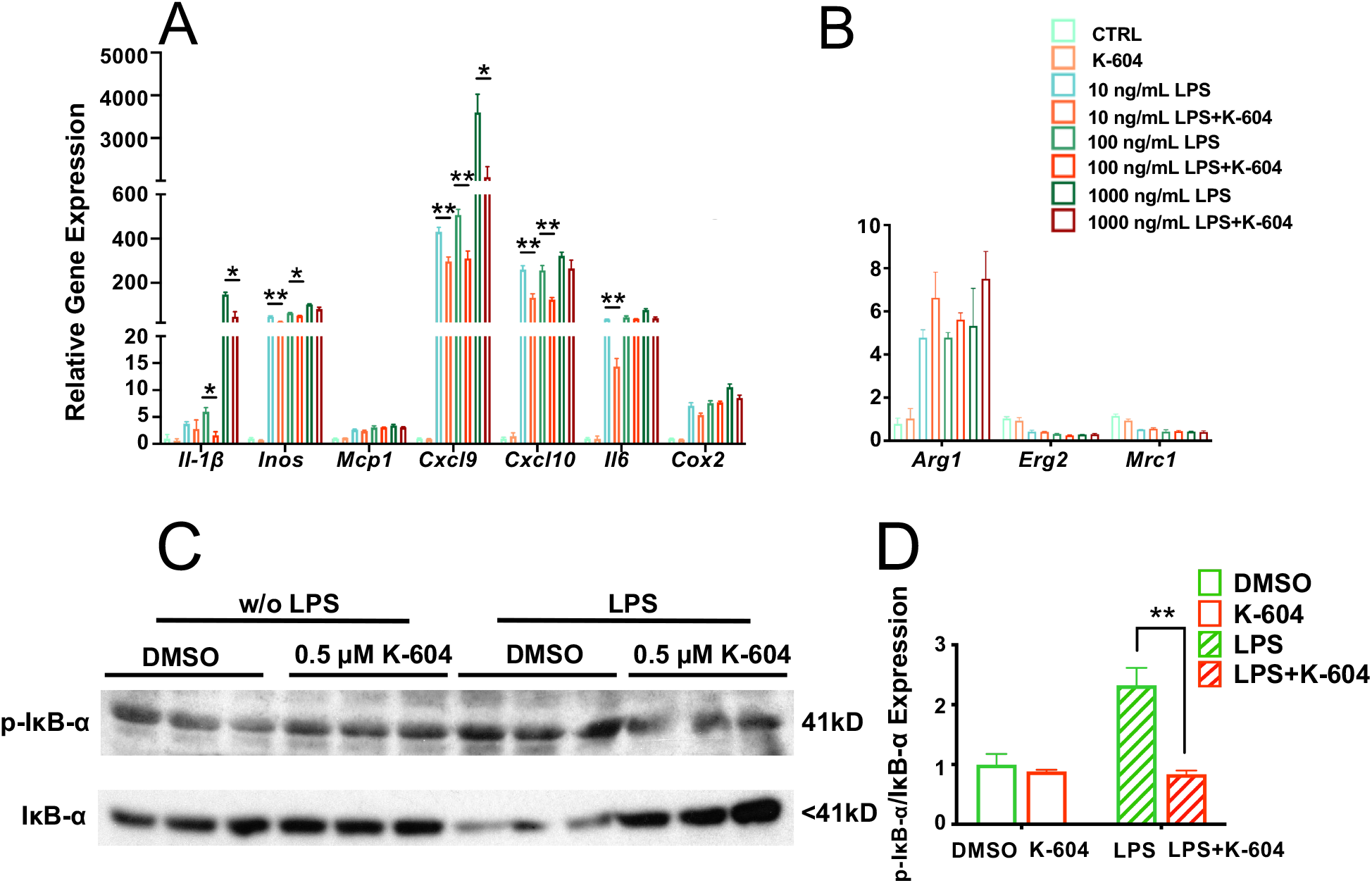
Inhibiting ACAT1 in N9 microglial cell line ameliorates proinflammatory responses towards LPS. N9 microglial cells were seeded at 1×10^5^ cells per well onto 12-well plates in medium RPMI-1640 with 10% calf serum. **A, B**, N9 cells were treated with DMSO (control group) or 0.5 μM ACAT1 inhibitor K-604 for four hrs, then treated with 10, 100, or 1000 ng/mL LPS as indicated. 6 hrs later, mRNAs were extracted, and real-time PCR was performed for pro-inflammatory (**A**) or anti-inflammatory (**B**) gene expressions. (**C), (D)**, N9 cells were treated with DMSO (control group) or 0.5 μM K-604 for 4 hrs, then treated with or without 100 ng/mL LPS for 30 min. Whole cell proteins were used for Western Blot to monitor phosphorylation status of IκB-α according to Methods described in Experimental Procedures.

In cells of the myeloid lineage, the TLR4 plays a major role in mediating pro-inflammatory cascades [37,38]. Activation of TLR4 by LPS or other TLR4 agonists triggers two signaling cascades [23]: the first involves TLR4 and the adaptor proteins TIRAP and MyD88 at the plasma membrane (PM) [24]; this MyD88-dependent signaling is terminated by endocytosis of TLR4 from the PM. The second TLR4 signaling cascade occurs non-canonically, after TLR4 is endocytosed, and involves other adaptor proteins TRAM and TRIF at the endosomes. After endocytosis, TLR4 cycles back to the PM, or eventually enters lysosomal compartment to be degraded [24]. Both signaling events involve activating the IκB kinase in the cytosol, which rapidly phosphorylates IκB and leads to its degradation (by the proteasome), and causes the transcription factor NFκB to move from the cytosol into the nucleus to activate the transcription of various pro-inflammatory genes [25]. To test if A1B inhibits LPS-dependent IκB kinase signaling, we treated N9 cells with or without K-604 for 4 hrs, then acutely exposed cells to LPS for 30 min, and used the cell extracts to monitor the phosphorylation status of IκB by Western blot analyses. The results show that, LPS treatment rapidly increased the phosphorylation status of IκB; pre-treating cells with A1B significantly abolished the effect of LPS on phosphorylation status of IκB, and significantly slowed down its degradation **(Fig. 3C, D)**. These results indicate that in N9 cells, A1B attenuates LPS-mediated inflammation in part by suppressing the ability of LPS to activate the IκB kinase.

### A1B alters the intracellular distribution of TLR4 and increases endocytosis of TLR4 in N9 cells treated with LPS for 30 min

At the PM, TLR4 undergoes endocytosis, membrane recycling that involve Golgi, endosomes, and PM, and eventually enters the lysosomal compartment for degradation. The rate of TLR4 endocytosis from the PM affects the LPS-induced pro-inflammatory responses {Reviewed in [23]}. Endocytosis of TLR4 can occur through clathrin-mediated process [26] or through caveolae/lipid rafts mediated process [27]. It is known that after LPS stimulation, a portion of TLR4 is enriched at the caveolae/lipid rafts region of the PM [27]. A1B causes cholesterol translocation from the endoplasmic reticulum where ACAT1 resides, to various membrane organelles, including the PM and various other membrane organelles [43, 44]. In the current cell system, we suspect that pre-treating cells with A1B may affect the intracellular distribution of TLR4, especially upon LPS stimulation. To test these possibilities, we first pretreated N9 cells with DMSO (control group) or with 0.5 μM K-604 for 4 hrs, then exposed them with or without LPS at 200 ng/ml for a very short period (30 min), the timeline of the experiment was shown in **Fig 4A**. We then prepared whole cell homogenates and subjected the homogenates to biochemical fractionation by OptiPrep ultracentrifugation.

**Fig 4.**
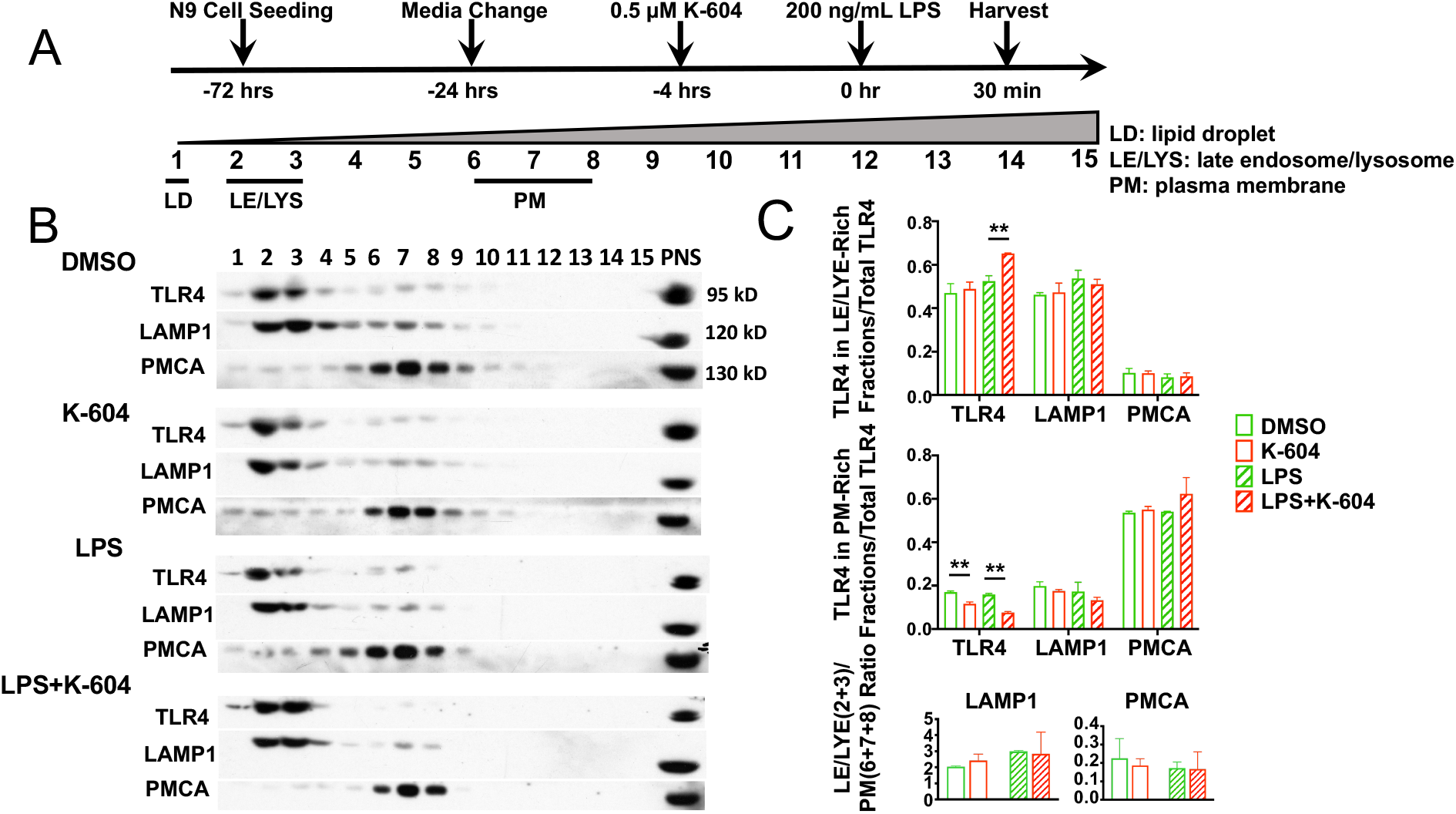
ACAT1 inhibition by K604 causes TLR4 enrichment in late endosome/lysosome in N9 cells acutely treated with LPS. **A**. Experimental design, and organelle marker distribution patterns after OptiPrep™ density gradient ultracentrifugation. Cells were treated with or without LPS from 30 min. **D**, fractions were analyzed for TLR4 content as well as for late endosome/lysosome marker LAMP1 and plasma membrane marker PMCA (Plasma membrane-type Ca^2+^-ATPases). The ratio of late endosome/lysosome fractions (2+3) to plasma membrane fractions (6+7+8) was used to determine their distribution between LE/LYS and PM. Calculation was made based on results from 3 individual experiments. Western blot analyses shown were from one of these experiments.

This procedure produces partial separation of various membrane organelles, primarily based on their differences in buoyant densities [28]. Here we subjected the individual OptiPrep fractions (1-15) by Western blot to monitor the distribution of TLR4. LAMP1 and plasma membrane calcium ATPase (PMCA) were used as the late endo/lysosome(LE/LYS) and PM markers, respectively. The results showed that, TLR4 is mainly distributed in 2 separate fractions: LE/LYS (fractions 2-3) and PM (fractions 6-8), with the contents in the LE/LYS fraction dominating over those in the PM fraction. Treating cells with A1B followed by LPS treatment for 30 min significantly increased the content of TLR in the LE/LYS fractions (**Fig 4C**, upper panel). In addition, treating cells with A1B decreased TLR4 by 50% in the PM, both with and without LPS (**Fig 4C**, middle panel). The protein levels of marker proteins LAMP1 and PMCA at the LE/LYS fraction vs those at the PM didn’t change (**Fig 4C**, lower panel). These results show that pretreatment with A1B causes more TLR4 to accumulate in the LE/LYS fractions, especially under acute LPS treatment.

The results described in **Fig 4** suggest that A1B increases the endocytosis rate of TLR4, especially upon acute LPS treatment. To test this hypothesis by using a direct assay, we performed an endocytosis assay in intact N9 cells based on the protocol described in [18]. Cells were pretreated with or without A1B, next treated with or without LPS for 30 min, then undergo endocytosis assay, by feeding cells with specific antibodies against TLR4. After endocytosis of the TLR/antibodies complex, cell surface-bound antibodies were removed; cells were then fixed, permeabilized with 0.3% Triton, stained with fluorescent secondary antibodies, and viewed and quantitated as the mean fluorescent area per cell under a confocal microscopy. The results (**Fig 5**) show that in cells not treated with LPS, pre-treatment of K-604 increased the amount of internalized TLR4 by around 45%; LPS treatment increased the amount of internalized TLR4 by 81%; pre-treatment of K-604 followed by LPS further increased the amount of internalized TLR4 by 33%. These results support the interpretation that A1B acts by increasing the endocytosis of TLR4 from the PM to the cell interior, especially in cells acutely treated with LPS.

**Fig 5.**
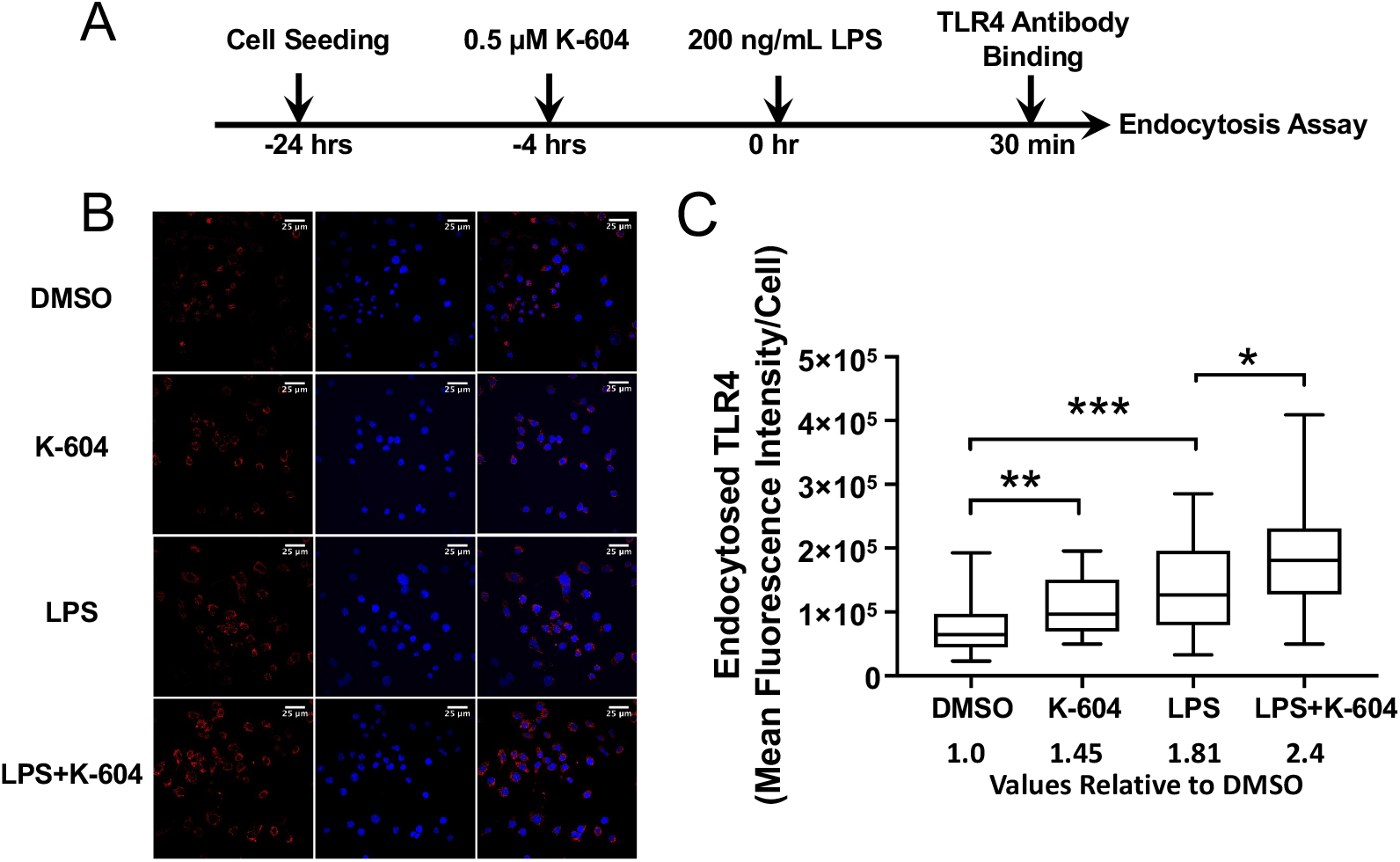
ACAT1 inhibition by K604 increases endocytosis of TLR4 in N9 cells. N9 cells seeded on poly-d-lysine-coated glass cover slides in 12-well plates were treated with DMSO or 0.5 μM K-604 for 4 hrs followed by treating with or without 200 ng/mL LPS for very short period (30 min). Endocytosis assay was performed as described in Methods. After treating cells with the primary antibodies against TLR4, the secondary antibodies (red) were used to stain TLR in the cell interior; DAPI stain (blue) was used to stain the nuclei. Quantification shows the mean fluorescent TLR signal per cell. The TLR signals were normalized by the nucleus signals. Calculation was made based on results from cells of 3 individual slides. Images shown were from one of these slides.

### A1B decreases TLR4 protein content in microglial N9 cells chronically treated with LPS (for 24-48 hrs)

The results described in **Figs 3–5** address the effect of A1B on TLR4 in N9 cells acutely treated with or without LPS for 30 min. Prolonged accumulation of TLR4 in the endosomes may lead to more TLR4 to enter the lysosomes and cause it to be degraded [26]. We tested this possibility by monitoring the effects of A1B on TLR4 in N9 cells treated with or without LPS for much longer period (24 hr or 48 hr). We first used double immunofluorescence staining in fixed permeabilized cells to monitor TLR4 content and cellular distribution; the PM protein marker N-cadherin was used as the control. The timeline of experiment is shown in **Fig 6A i**. The results (**Fig 6A ii**) show that, without LPS, A1B does not alter the cellular content of TLR4 protein. In contrast, in cells treated with LPS for 24 hrs, A1B caused a significant decrease (by 28%) of the TLR4 protein levels. Additional analyses of the imaging result (**Fig 6A iv**) show that either with or without LPS treatment, A1B does not alter the relative distribution of TLR4 between the internal membrane (IM) and the PM. This result suggests that pre-treating N9 cells with A1B causes reduction of TLR4, when cells are treated chronically with LPS. To strengthen this finding by using a different method, we repeated the experiment by using the same condition as described in **Fig 6A** (but plated the cells with higher initial cell density). At different time points, we prepared homogenates from cells to determine the total TLR4 contents by Western blot analysis. Vinculin was used as the loading control. The results (**Fig 6B ii, iii**) show that in cells with or without LPS treatment for 24 hrs, A1B by K-604 does not alter the TLR4 protein content. In contrast, in cells treated with LPS for 48 hrs, A1B caused a significant decrease (by 25%) of the TLR4 protein. This result combined with the results shown in **Figs 4** and **5** show that in cells acutely treated with LPS, A1B causes TLR4 at the PM to be endocytosed at a more rapid rate. In N9 cells chronically treated with LPS for 24-48 hrs, A1B causes more TLR4 to be degraded, perhaps as a consequence of more TLR4 entering the late endo/lysosomes. The results described above show that A1B alters the fate of TLR4 in LPS treated microglial N9 cells, by increasing its endocytosis and by decreasing its total protein content. Overall, these results obtained in cell culture (**Figs 3–6**) support the notion that blocking ACAT1 in microglia suppress the ability of TLR4 in pro-inflammatory signaling in response to LPS by modulating its intracellular fate.

**Fig 6.**
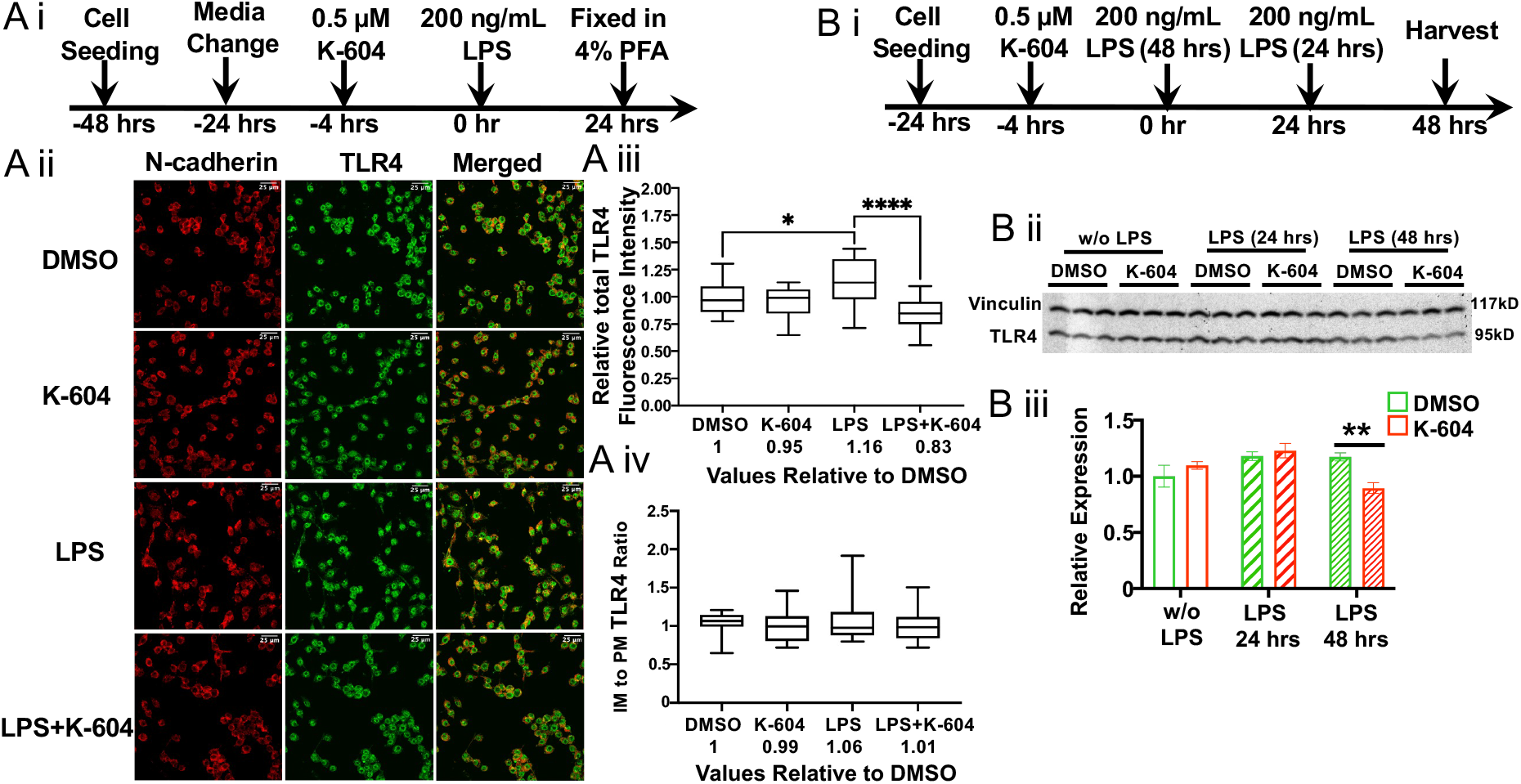
A1B decreases TLR4 protein content in microglia chronically treated with LPS. N9 cells seeded at 1×10^5^ cells per well on poly-d-lysine-coated glass cover slides in 6-well plates were pre-treated 4-hrs with DMSO or with 0.5 μM K-604, then exposed with or without 200 ng/mL LPS for 24-hrs. Double immunofluorescence staining for TLR4 and for the plasma membrane marker N-cadherin was then performed. **I**. Timeline of the experiment. II. Representative images demonstrating TLR4 distribution in N9 cells; **III**. Quantification of total TLR4 relative fluorescence intensity per cell; **IV**. Quantification of IM/PM TLR4 fluorescence intensity ratio. 15 cells per group were analyzed. The TLR4 signals overlapping with that of N-cadherin are considered as TLR4 at the PM, while those not overlapping are considered as TLR4 at the IM. N9 microglial cells were seeded at 2×10^5^ cells per well onto 6-well plates in RPMI-1640 with 10% calf serum. N9 cells were treated with DMSO (control group) or 0.5 μM K-604 for 4 hrs then treated with or without 200 ng/mL LPS for 24 and 48 hrs. At different time points, cells were harvested for protein isolation and TLR4 Western Blot analyses. Vinculin was used as the protein loading control. I. Timeline of experiment. II. Western blot. III. Quantitation of Western blot.

### A working hypothesis to explain the A1B actions on suppressing LPS-mediated pro-inflammatory signaling cascade in microglia (Fig 7)

Activation of TLR4 by LPS triggers two signaling cascades [23]: the first involves TLR4 and the adaptor proteins TIRAP and MyD88 at the PM; this signaling process is terminated by endocytosis of TLR4 from the PM. After TLR4 is endocytosed, the second TLR4 signaling cascade occurs non-canonically at the endosomes and involves other adaptor proteins TRAM and TRIF. Regarding the first signaling cascade: LPS binds to CD14 or other coreceptors such as CD36 and causes the TLR4/LPS complex to move laterally to be concentrated at certain cholesterol-rich lipid raft microdomain at the PM, to participate in pro-inflammatory signaling, and to undergo caveolae-mediated endocytosis [27]. We hypothesize that pre-treating cells with A1B causes subtle change(s) of the cholesterol-rich microdomain at the PM such that the TLR4/LPS complex enriched at the lipid raft domain is endocytosed at a more rapid rate, causing the first signaling event to be terminated more effectively. We also hypothesize that in cells treated with LPS for longer time (24-48 hrs), A1B causes more TLR4 to accumulate in the lysosomal compartment, causing it to be degraded, thereby depleting the total TLR4 content. After TLR4 is endocytosed, the second TLR4 signaling cascade occurs non-canonically at the endosomes and involves other adaptor proteins TRAM and TRIF. A1B may or may not affect clathrin-mediated endocytosis of TLR4, and/or affect the second TLR4 signaling cascade.

**Fig 7.**
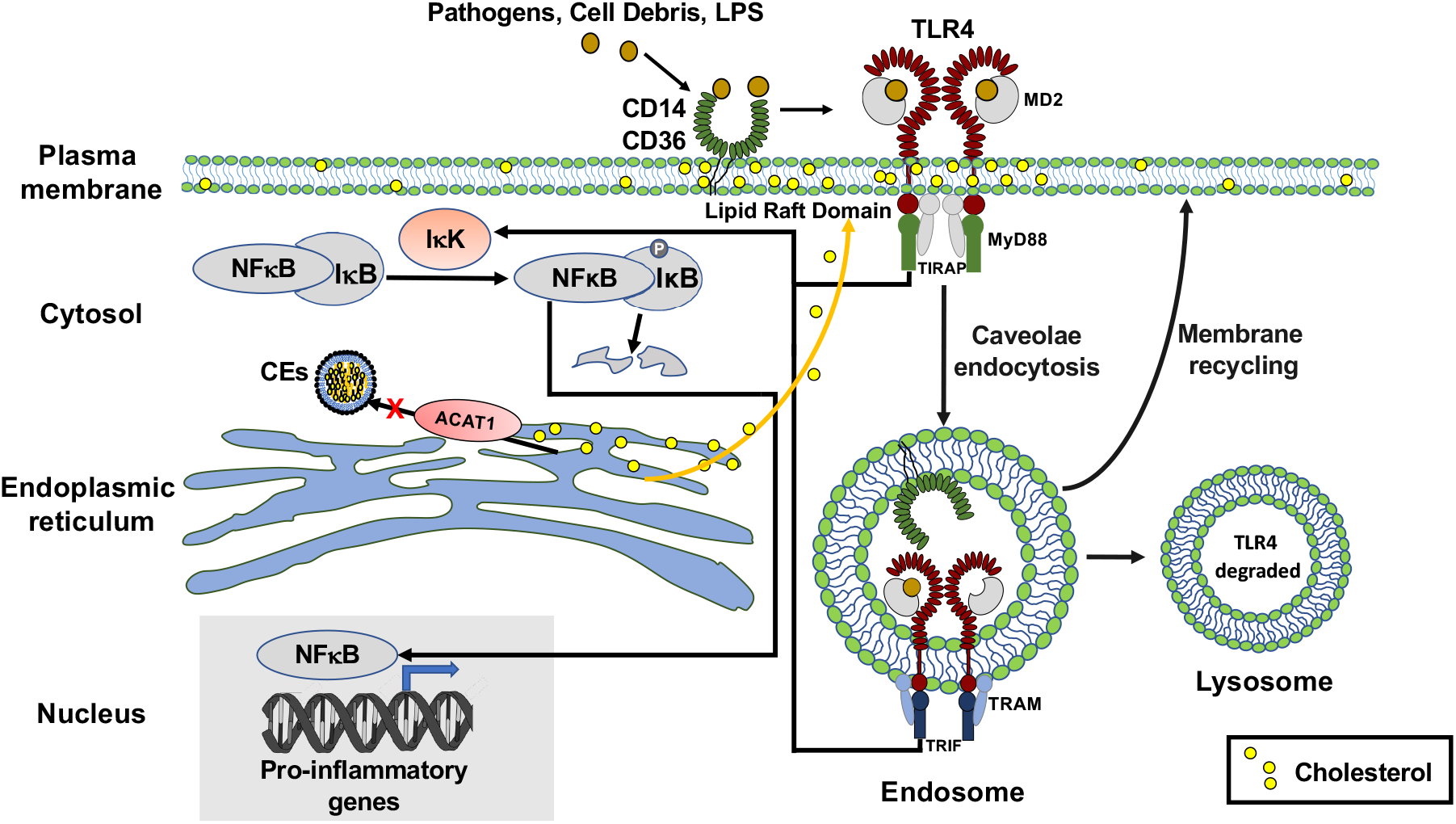
A working model to account for the action of A1B to suppress proinflammatory signaling cascade in microglia treated with LPS. At the PM of microglia, LPS (or other pathogens/oxidized lipids) bind to TLR4/MD2 complex (with CD14 or CD36 as co-receptors) to trigger the dimerization of TLR4/MD2 within the lipid raft domain. The adaptor proteins MyD88/TIRAP bound to TLR4/MD2 activates IKK mediated phosphorylation of IκB, causing nuclear translocation of NFκB to activate transcriptions of various proinflammatory genes. TLR4/MD2 undergoes caveolae/lipid raft mediated endocytosis, this process terminates its proinflammatory signaling activities at the PM. ACAT1 is located at the endoplasmic reticulum. In cells acutely treated with LPS, A1B prevents the cholesterol pool from forming cholesterol esters (CEs) such that it translocated to the lipid raft domain at the PM and causes TLR4 to undergo caveolae mediated endocytosis at more rapid rate. TLR4/MD2 complex at the endosome can initiate a second proinflammatory signaling event that involves similar but different adapter proteins (TRAM, TRIF; etc.). Whether A1B also affects the signaling activity of TLR4/MD2 at endosomes is currently unknown. TLR4/MD2 complex at the endosome undergoes membrane recycling to return to the PM; it eventually enters the terminal lysosomal compartment to be degraded. In N9 cells, chronically treated with LPS, A1B may act by increasing the residence time of TLR4 in the lysosomes, such that more TLR4 gets degraded.

## Discussion

ACATs/SOATs are potential targets to treat several human diseases, including atherosclerosis [29], certain forms of cancer {early literature reviewed in [5]}, [30], and neurodegenerative diseases that include AD [31] and NPCD [32]. There are two *Acats/Soats* in mammals [4]. To link ACATs with neurodegeneration, it is important to document the relative expressions of these two genes in the CNS. In the current work, we retrieved data from literature to demonstrate that in the CNS: **1**. under non-disease condition, in the human brain and the mouse brain, the expression of *Acat1/Soat1* dominates over that of *Acat1/Soat2* (**Fig S1**); **2**. *Acat1/Soat1* is expressed in various cell types, with microglia expressing the highest level (**Fig S1**); **3**. the expression of *Acat1/Soat1* in microglia are significantly elevated in mouse models for acute inflammation by LPS, for chronic inflammation in various neurodegenerative diseases, and for aging; **4**. its expression is also significantly elevated in vulnerable (entorhinal cortex) regions of patients with LOAD (**Fig S2**). It is known that under these conditions, and in the hippocampus and other vulnerable regions of the LOAD patient brain, the levels of the pro-inflammatory cytokine TNF-α are significantly increased [33, 34]. In human monocytes/macrophages, the pro-inflammatory cytokine TNF-α significantly increased *ACAT1/SOAT1* mRNA and ACAT1 protein levels [35]. It is possible that, in both acute and chronic inflammatory contexts, increases in TNF-α and other pro-inflammatory cytokines play a key role in upregulating *SOAT1* [36, 37]. The validity of this hypothesis needs to be tested by future experimentation.

Current results show that cell-type specific A1B (by genetic inactivation of *Acat1/Soat1*) in microglia, but not in neurons, significantly attenuated the pro-inflammatory responses induced by *LPS in vivo* (**Figs 1** and **2**). To pursue the mechanism(s) of action of A1B, we used the mouse microglial N9 cells and treated the cells with K-604. The results show that: in cells acutely treated with LPS (for 30 min), pre-incubating cells with A1B significantly increased the endocytosis of TLR4, the major receptor for mediating the LPS initiated signaling at the PM (**Figs 4** and **5**); in cells chronically treated with LPS (for 24-48 hrs), pre-incubating cells with A1B significantly decreased the total TLR4 protein content, presumably as a consequence of enhanced trafficking of TLR4 to the lysosomes, which is the degradative compartment for many cellular proteins. Overall, our results show that A1B suppresses LPS-mediated neuroinflammation at least in part by modulating TLR4-mediated signaling. In various immune cells, in addition to TLR4, several other TLRs also play important roles in mediating pro-inflammatory signaling cascade events. Our results cannot rule out the possibility that A1B suppresses LPS-induced pro-inflammatory responses by additional mechanism(s) that are independent of TLR4. We had previously reported that blocking ACAT1 in macrophages ameliorates pro-inflammatory responses caused by LPS [12] or by cholesterol loading [13]. It is known that in macrophages, TLR4 plays important roles in mediating various chronic pro-inflammatory signaling initiated by oxidized low-density lipoprotein [36]. Based on our current results, A1B may suppress oxidized low-density lipoprotein mediated inflammation by modulating the fate of TLR4. This possibility needs to be investigated further.

At present, the mechanism by which A1B can cause increase in endocytosis of TLR4 is unclear. It has been shown that upon activation by various ligands, TLR4 (and other TLRs) at the PM as monomer undergoes lateral diffusion, dimerizes, and becomes enriched at a certain cholesterol-rich/sphingolipid-rich microdomain (i.e. lipid rafts/caveolae) at the PM [38]. We have recently reported that in mouse embryonic fibroblast cells, A1B causes redistribution of cholesterol contents in multiple cellular compartments [32]. Thus, A1B could alter the cholesterol content of the TLR4 associated membrane microdomain at the PM and causes this domain to undergo caveolae-mediated endocytoses at a more rapid rate. Other possibilities also exist. For example, in addition to affecting its endocytosis rate, A1B may also affect the rates of recycling of TLR4 containing membranes back to the PM, and/or its entry to the lysosomes. These and other possibilities will need to be further investigated.

In CNS, TLR4 is expressed in microglia, astrocytes, oligodendrocytes, and neurons. TLR4 recognizes diverse pathogen-derived ligands including LPS, and various tissue damage-related ligands including oligomeric amyloid peptide fragment Aβ_1-42_, heat-shock proteins (especially HSP60 and HSP70), high mobility group box 1 (HMGB1); etc. It plays a key role in mediating pro-inflammatory responses caused by various infectious and non-infectious agents, and is a LOAD susceptibility gene [39]. It will be interesting to examine if A1B suppresses chronic inflammation that occurs in neurodegenerative diseases including AD and related dementias, by modulating the cellular fate of TLR4.

## Abbreviations

Aβ: amyloid beta peptides
ACAT1/SOAT1: acyl-coenzyme A:cholesterol acyltransferase 1/sterol O-acyltransferase 1
AD: Alzheimer’s disease
CEs: cholesteryl esters
CNS: central nervous system
EOAD: early onset Alzheimer’s disease
ER: endoplasmic reticulum
hAPP: human amyloid precursor protein
IM: internal membranes
iPSC: Induced pluripotent stem cell
KO: knock-out
LE: late endosome
LOAD: late onset Alzheimer’s disease
LPS: lipopolysaccharides
Lys: lysosomes
PM: plasma membrane
SEM: standard error of the mean
Tg: transgenic
TLR4: toll-like receptor 4.
TNF_alpha_: tumor necrosis factor alpha
TREM-2: Triggering Receptor Expressed On Myeloid Cells 2

## Supplementary Information

Fig S1 and Fig S2.

## Acknowledgements

We thank Ann Lavanway and Zdenek Svindrych, for advice on confocal microscopy usage; members of the Chang laboratory for helpful discussions during the course of this work; and Dr. Gustav Lienhard for critical reading of this manuscript.

## Author contributions

H.L., T.H., C.C.Y.C. and T.-Y.C. designed research; H.L., T.H., M.D. and J.G. performed research; H.L., T.H., M.D. analyzed data; T.-Y.C., C.C.Y.C., J.G., and T.H. wrote and edit the manuscript.

## Funding

This work was supported by NIH Grant R01 AG063544 (to T.-Y.C and C.C.Y.C.). We acknowledge the shared facilities of the preclinical Imaging and Microscopy Resource and National Cancer Institute Cancer Center Support Grant 5P30 CA023108-37 at the Norris Cotton Cancer Center at Dartmouth, and NIH Grant P20-GM113132 to support the Institute for Biomolecular Targeting at Dartmouth. Additional funding was provided by a National Institute on Aging fellowship F30 AG074524 (to M.T.D.).

## Declarations

### Ethics approval and consent to participate

All the experiments were performed in accordance with legal and institutional guidelines and were carried out under ethics, consent and permissions of the Ethical Committee of Care and Use of Laboratory Animals at Geisel School of Medicine at Dartmouth.

### Consent for publication

Not applicable.

### Availability of data and materials

The authors declares that the relevant data are included in the article.

### Competing Interests

The authors declared that they are no competing interests.

---

**Fig S1.**
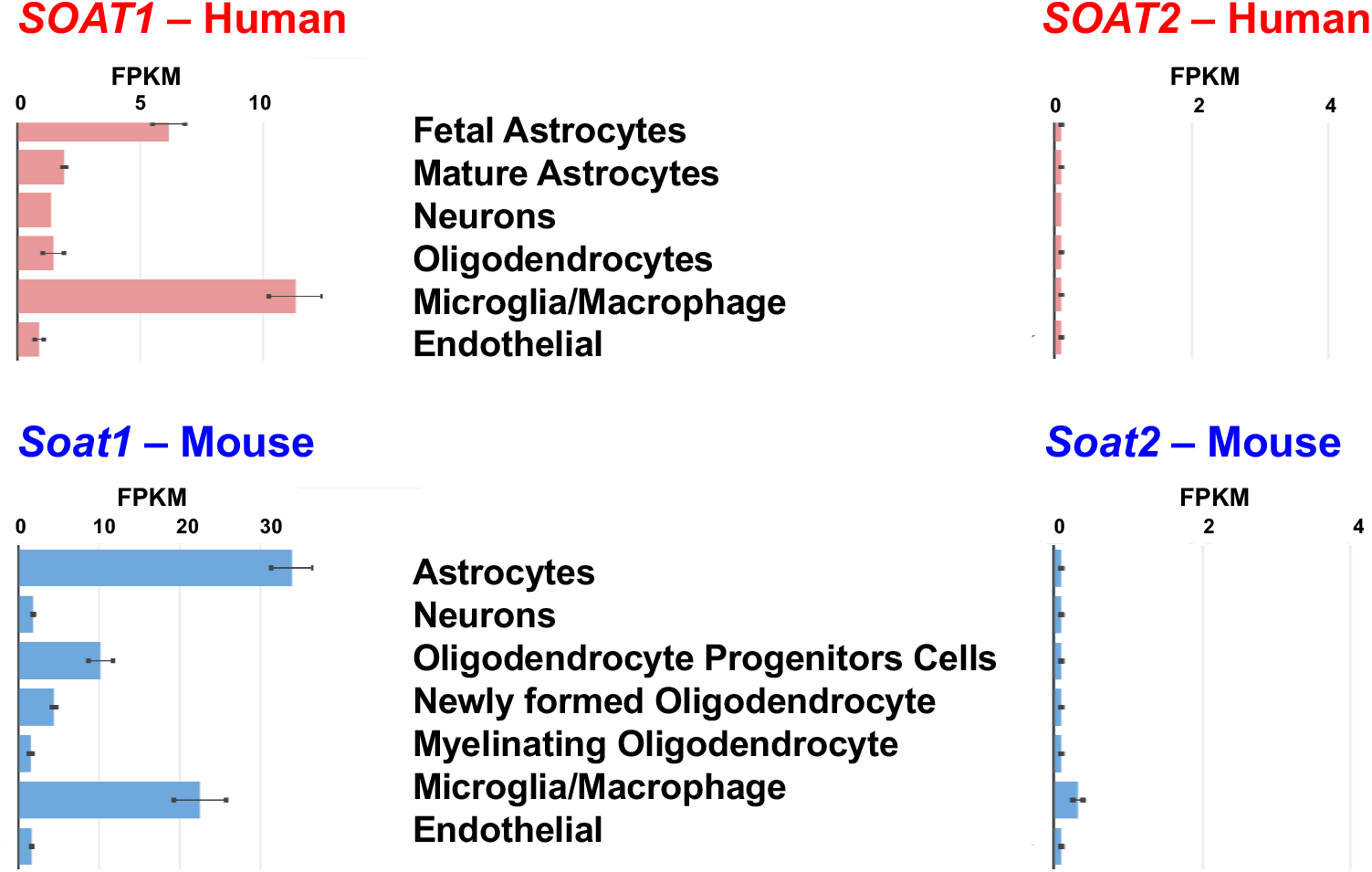
*ACAT1/SOAT1, but not ACAT2/SOAT2*, is significantly expressed in both human and mouse microglia. Microglia express high levels of (A) *ACAT1/SOAT1* in human brains and (B) *Acat1/Soat1* in mouse brains. Gene expression data are in fragments per kilobase million (FPKM). Data are retrieved from the Barres Laboratory [23]. Error bars represent +/− 1 SEM.

**Fig S2.**
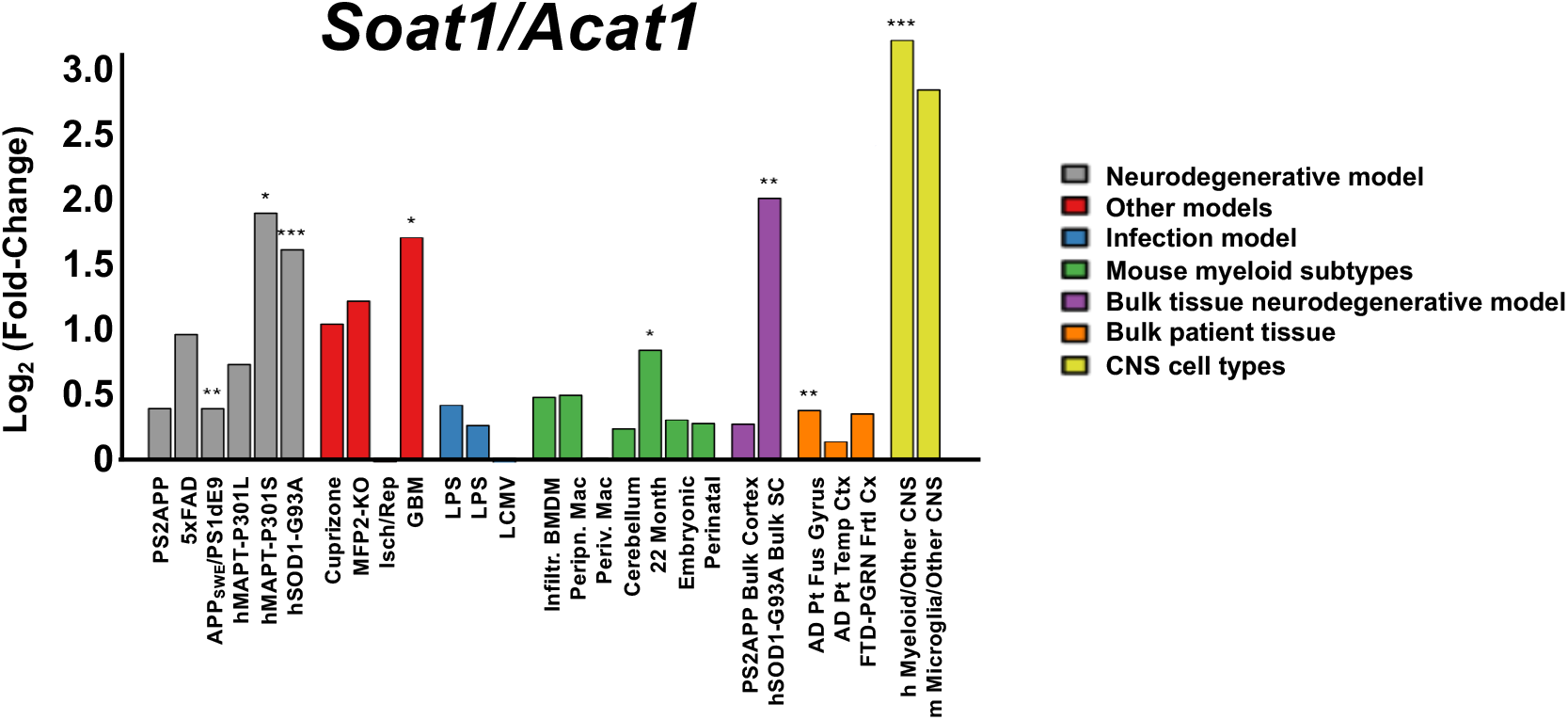
*Acat1/Soat1* expression is up-regulated in several human patient brains and mouse models of neurological diseases. Mouse model expression includes microglia from PSEN2/APP, 5XFAD, APP(Swedish)/PSEN1(dE9), MAPT(P301L), MAPT(P301S), and SOD1(G93A). Human AD patient tissue includes fusiform gyrus and temporal cortex. Gene expression data are log2 (fold-change). Data are retrieved from Friedman and colleagues [24].

## Notes

### Competing Interest Statement

The authors have declared no competing interest.

